# DNA replication initiation in *Bacillus subtilis*; Structural and functional characterisation of the essential DnaA-DnaD interaction

**DOI:** 10.1101/444885

**Authors:** Eleyna Martin, Huw E. L. Williams, Matthaios Pitoulias, Daniel Stevens, Charles Winterhalter, Timothy D. Craggs, Heath Murray, Mark S. Searle, Panos Soultanas

**Affiliations:** Centre for Biomolecular Sciences, School of Chemistry, University of Nottingham, Nottingham NG7 2RD, UK; Centre for Bacterial Cell Biology, Medical School, Newcastle University, Newcastle NE2 4AX, UK; School of Chemistry, University of Sheffield, Western Bank, Sheffield S10 2TN UK

**Keywords:** *Bacillus subtilis*, DNA replication, protein NMR, protein-protein interactions, DnaD, DnaA

## Abstract

The homotetrameric DnaD protein is essential in low G+C content gram positive bacteria and is involved in replication initiation at *oriC* and re-start of collapsed replication forks. It interacts with the ubiquitously conserved bacterial master replication initiation protein DnaA at the *oriC* but structural and functional details of this interaction are lacking, thus contributing to our incomplete understanding of the molecular details that underpin replication initiation in bacteria. DnaD comprises N-terminal (DDBH1) and C-terminal (DDBH2) domains, with contradicting bacterial two-hybrid and yeast two-hybrid studies suggesting that either the former or the latter interact with DnaA, respectively. Using Nuclear Magnetic Resonance (NMR) we show that both DDBH1 and DDBH2 interact with the N-terminal domain I of DnaA and studied the DDBH2 interaction in structural detail by NMR. We revealed two families of conformations for the DDBH2-DnaA domain I complex and showed that the DnaA-interaction patch of DnaD is distinct from the DNA-interaction patch, suggesting that DnaD can bind simultaneously DNA and DnaA. Using sensitive single-molecule FRET techniques we revealed that DnaD remodels DnaA-DNA filaments consistent with stretching and/or untwisting. Furthermore, the DNA binding activity of DnaD is redundant for this filament remodelling. This in turn suggests that DnaA and DnaD are working collaboratively in the *oriC* to locally melt the DNA duplex during replication initiation.

## Introduction

Replicating and propagating genomes is the raison d’être of all biological systems. The assembly of an “orisome” at dedicated genome sites, known as chromosomal origins (*oriC* in bacteria), is a carefully choreographed and regulated process involving a variety of proteins collectively known as replication initiators (**1-3**). Orisome assembly locally melts the *oriC* and facilitates loading of the replicative helicase and primase, the first and key step in the assembly of bidirectional replication forks (**4-6**). At a gross mechanistic level there are common features across all domains of life with replication initiation proteins phylogenetically related but molecular details have diverged considerably across different biological systems (**1, 3**).

Origin recognition complex (ORC) and Cdc6 proteins play crucial roles in eukaryotic replication initiation whereas the ubiquitous, strictly conserved AAA+ DnaA protein is the replication initiator in bacteria (**7-10**). DnaA comprises four domains (I-IV), with the C-terminal domain IV binding specific dsDNA sequences within the *oriC*, known as DnaA-boxes (**11**), and the central AAA+ (<u>A</u>TPases <u>A</u>ssociated with various cellular <u>A</u>ctivities) domain III forming right-handed oligomeric filaments that wrap dsDNA around the outside. These filamentous nucleoprotein assemblies impose positive toroidal strain on the *oriC* which is dissipated by unwinding and melting of the DUE (<u>D</u>NA <u>U</u>nwinding <u>E</u>lement) during replication initiation. DnaA filaments also invade and bind ssDNA generated during melting of the DUE, through a series of α-helices forming a ssDNA-binding staircase inside the central channel of the filament (**12, 13**). The role of the N-terminal domains I and II in the assembly of the initiation complex is the least understood (**14**). Domain II appears to be a rigid linker region of variable lengths in different bacterial species which connects domain I to the central AAA+ domain III whilst domain I interacts with different client proteins, like SirA involved in sporulation (**15**), HobA, Hda, DiaA and YabA involved in regulation of replication initiation (**16-19**) and the replicative helicase DnaB in *Escherichia coli* (**20, 21**).

Orisome assembly in low G+C-content *Firmicutes* requires two essential primosomal proteins DnaD and DnaB which are not found in *E. coli* and related bacteria. DnaD interacts with the *oriC*-DnaA complex and sequentially recruits DnaB and the DnaI-DnaC complex to load the replicative helicase DnaC at the *oriC* (**2**). Mutations in the corresponding *dnaD*, *dnaB* and *dnaI* genes causes defects in replication initiation and re-initiation (**22-24**). DnaD and DnaB are structural homologues sharing DDBH1 and DDBH2 (<u>D</u>naD <u>D</u>naB <u>H</u>omology 1 and 2, respectively) domains (**25**). They are thought to act together as co-loaders interacting with DnaI during loading of DnaC at the *oriC* (**26, 27**). DnaD comprises an N-terminal DDBH1 with a Winged-Helix fold (WH) forming tetramers that can further assemble into higher order oligomers, and a C-terminal DDBH2 which binds DNA with better affinity for ss than dsDNA (**28, 29**). Binding DDBH2 alone to dsDNA causes untwisting (partial melting) of the double helix which becomes more extensive when the full-length DnaD protein binds to dsDNA (**30, 31**), extending B-form duplex DNA from its normal 10.5 bp per helical turn to 16 bp per turn (**32**).

The formation of nucleoprotein structures by DnaA and DnaD and their interaction provide the foundation for orisome assembly and initiation of DNA replication in *Firmicutes*. Yet the molecular details that underpin this process are still unknown. Here, we used protein NMR to reveal structural details of the *Bacillus subtilis* DnaA and DnaD interaction. We show that both the N-terminal DDBH1 and the C-terminal DDBH2 of DnaD interact with the N-terminal domain I of DnaA. The latter complex is much weaker and modelling with NMR restraints reveals two distinct but overlapping conformations. Combined with biochemical and single-molecule FRET (<u>F</u>luorescence <u>R</u>esonance <u>E</u>nergy <u>T</u>ransfer) we also show that binding of DnaD to DnaA-DNA filaments induces conformational filament changes consistent with filament stretching/untwisting. The significance of this in terms of DnaD interacting with the DnaA-DnaD filament and orisome assembly is discussed.

## Results

### Identification of the interacting domains of DnaA and DnaD

Yeast-two hybrid studies revealed that deleting the C-terminal part of DnaD, residues 140-232, had no effect on its interaction with DnaA, but truncating the protein for a further seven residues to amino acid residue 133 abolished the interaction (**33**). This patch of seven amino acid residues is located at the N-terminal end of the C-terminal half of the DnaD protein (residues 129-232) also found in the DDBH2 (**25**). However, a recent bacterial two hybrid study contradicted this and showed that the DnaD DDBH1 (residues 1-128) and not the DDBH2 interacts with the N-terminal domain I (residues 1-82) of DnaA (**34**). The N-terminal domain I of DnaA also interacts with the replicative helicase (**20, 21**) and other client proteins involved in the regulation of replication initiation (**16-19**) and as part of a regulatory interaction hub, it may also be involved in the interaction with DnaD. In order to clarify this contradiction we cloned, recombinantly expressed and purified the domain I of DnaA and the DDBH1 and DDBH2 domains of DnaD in order to investigate their interactions and their significance (**Suppl. Fig. S1**).

As the DnaD DDBH1 has recently been reported to interact strongly with the DnaA domain I (**34**) we first tested this interaction by NMR. An HSQC titration of DnaD DDBH1 into ^15^N-labelled DnaA domain I revealed significant broadening of the DnaA domain I backbone NH signals in a concentration dependent manner and recovery of the signal was not observed by a ratio of DnaA domain I:DnaD DDBH1 of 1:4 (**Fig. 1A**). This is strongly indicative of an interaction but extensive line broadening precluded a detailed NMR study.

**Figure 1.**
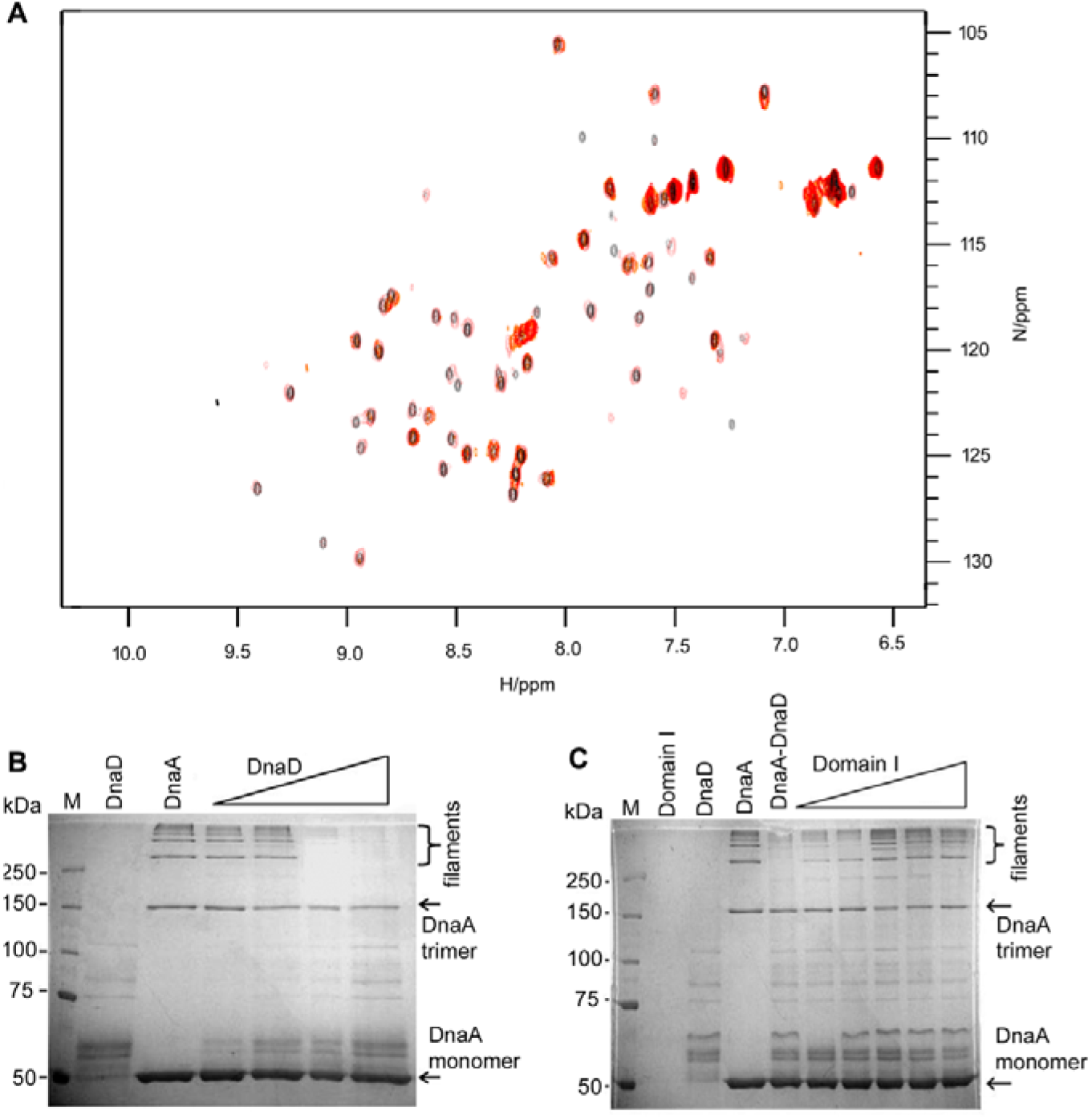
**A.** HSQC Titration of DnaD-DDBH1 domain into ^15^N-labelled DnaA domain I (100 μM), showing ratios (DnaA-DDBH1:DnaD domain I) of 1:0 (Grey), 1:0.27 (pink) 1:0.75 (orange) 1:1.75 (red). Data show extensive broadening of all peaks with a small number of sidechain NH protons remaining visible at ratios greater than 1.75. **B.** Analysis of DnaA(N191C/A198C)-DNA filaments (DnaA(N191C/A198C) 1.5 μM, DNA 3 nM) incubated with increasing concentrations of DnaD (1.5, 3, 6, and 12 μM), shown by the triangle. Lane M shows molecular weight markers, as indicated, and the rest of the lanes from left to right show DnaD (12 μM) on its own (too small to see as it has run off the bottom of the gel, the visible bands are impurities carried over from the purification of untagged DnaD) and DnaA(N191C/A198C)-DNA filaments formed in the absence of DnaD. The gel shows clearly that the formation of DnaA(N191C/A198C)-DNA filaments is inhibited with increasing concentrations of DnaD. **C.** The same experiment as that shown above was repeated but this time the DnaA(N191C/A198C)-DNA filaments were incubated with 6μM DnaD and increasing concentrations of DnaA domain I (1.5, 3, 6 and 12 μM). Lane M shows molecular weight markers, as indicated, and the rest of the lanes from left to right show DnaA domain I (12 μM) (too small to see as it has run off the bottom of the gel), DnaD (12 μM) and DnaA(N191C/A198C)-DnaD filaments forming in the absence of DnaD and DnaA domain I.

In order to confirm that it is the domain I of DnaA that interacts with DnaD we utilised DnaA-DNA filaments which have been previously detected *in vivo* (**35**) and *in vitro* (**36**). In the latter case, cysteines were introduced at positions N191 and A198, and utilised to chemically cross-link neighbouring molecules within the DnaA-DNA filament using BMOE (bis-maleimido ethane). Cross-linked DnaA species were then resolved by SDS-PAGE and visualised by western blotting. The interaction of DnaD with the DnaA-DNA filaments abolished BMOE cross-linking of neighbouring DnaA molecules within the filament which was detected by the apparent disappearance of higher ordered cross-linked species. We argued that if domain I of DnaA interacts with DnaD it should then be able to sequester DnaD in this assay and alleviate the apparent inhibitory effect of DnaD on the chemical cross-linking of DnaA molecules.

To test this, we constructed the DnaA(N191C/A198C) mutant protein and repeated the BMOE cross-linking assays in the presence of increasing concentrations of DnaD and an 819 bp DNA fragment containing the half origin with the DUE and four DnaA-boxes between the *dnaA* and *dnaN* genes (**Suppl. Fig. S2 and S3**). Higher order DnaA filaments induced by binding to DNA were clearly visible which disappeared at higher concentrations of DnaD (**Fig. 1B**). Increasing concentrations of DnaA domain I sequestered DnaD and restored the appearance of higher order DnaA-DNA filaments, indirectly suggesting that it is the domain I of DnaA that interacts with DnaD (**Fig. 1C**). In order to investigate whether the DnaD DDBH2 also interacts with the DnaA domain I and precisely map the interaction interfaces of the two proteins we utilised protein NMR techniques.

### NMR analysis of the N-terminal DnaA domain I and mapping of the interaction interface with DnaD

The DnaA domain I (residues 1-81) was ^13^C/^15^N isotopically labelled and a range of 2D and 3D heteronuclear NMR experiments at 800 MHz (see methods) allowed 97% of the non-prolyl residues of the DnaA domain I amide backbone to be assigned. The detailed backbone assignment provided the basis for mapping the interaction surface of DnaA using chemical shift perturbation (CSP) effects.

Unlabelled DnaD DDBH2 domain was titrated into a ^15^N-labelled sample of DnaA domain I (100 μM) up to an 8:1 excess and the binding interaction monitored incrementally in 2D ^1^H/^15^N HSQC spectra. Statistically significant perturbations for nine residues within the DnaA domain I were observed (**Fig. 2A and Suppl. Fig. S4**). The effects were mapped to the protein surface and found to correspond to a well-defined binding patch on one face of the domain, with residues clustered into three groups: Lys17, Ser20 and Ser23 located at the interface between the C-terminus of helix α1 and the N-terminus of helix α2, and Phe49, Ala50, Asp52, Trp53 and Glu55 located throughout helix α3, and His60 located at the N-terminus of helix α4 (**Fig. 2B**). The absence of any significant CSP effects within the β-sheet regions of the domain demonstrates that the interface involving DnaA domain I is extensively α-helical. The CSP effects observed during the titration were also used to generate binding isotherms and the data fitted to demonstrate a weak binding affinity (K_D_ = 768±168 μM) between DnaA domain I and the DnaD DDBH2 domain when averaged over 5 well resolved residues (S20, A50, D52, E55 and H60) (**Suppl. Fig. S4**).

**Figure 2.**
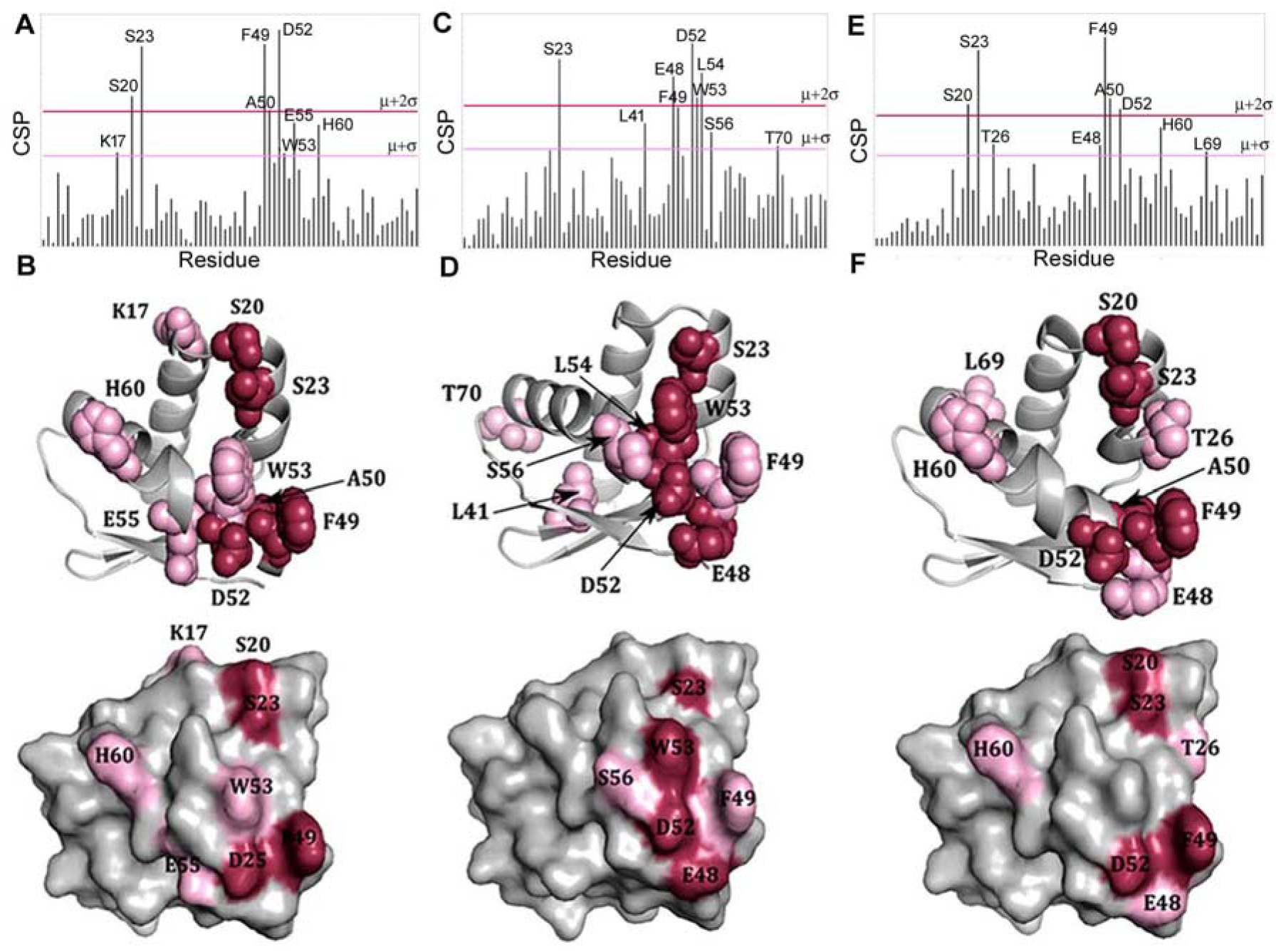
**A.** CSP analysis of the ^15^N-labelled DnaA domain I titration with the DnaD DDBH2 domain. Threshold values of significance were calculated using μ = 0.0139 and σ = 0.0134. The μ + 2σ and μ + σ limits are highlighted in red and pink respectively. **B.** Residues exceeding the threshold limits are mapped onto the X-ray crystal structure of DnaA domain I (pdb 4TPS). **C.** CSP analysis of the ^15^N-labelled DnaA domain I titration with the DnaD DDBH2 domain in the presence of ssDNA (800 μM). Threshold values of significance were calculated using μ = 0.0143 and σ = 0.0111. The μ + 2σ and μ + σ limits are highlighted in red and pink respectively. **D.** Residues exceeding the threshold limits are mapped onto the X-ray crystal structure of DnaA domain I (pdb 4TPS). **E.** CSP analysis of the ^15^N-labelled DnaA domain I titration with the DnaD DDBH2 domain truncation (residues 129 - 196). Threshold values of significance were calculated using μ = 0.0152 and σ = 0.0120. The μ + 2σ and μ + σ limits are highlighted in red and pink respectively. **F.** Residues exceeding the threshold limits are mapped onto the X-ray crystal structure of DnaA domain I (pdb 4TPS).

The DDBH2 domain of DnaD is involved in DNA binding (**29-31**). The conserved motif YxxxIxxxW (Y^180^IDRI^184^LFEW^188^) along with a region of the unstructured C-terminus (residues 206-215) appear to be essential for DNA binding (**25**). At the origin of replication, both proteins bind to each other and function to remodel the DNA for replication initiation (**2**). DnaD can bind both ds and ss DNA, but with a higher affinity for the latter. Consequently, a short 10-mer of ssDNA (5’-GTTATTGCTC) previously used in DnaD-DNA binding studies (**25**) was selected for studies of the tertiary interaction by NMR. We repeated the titration of unlabelled DnaD DDBH2 domain with ^15^N-labelled DnaA domain I under identical conditions, but in the presence of an 8-fold excess of ssDNA (800μM). Selective CSP effects mapped to a similar binding patch on DnaA with the cluster of residues Glu48, Phe49, Asp52, Trp53, Leu54, and Ser56, located on helix α3 helix, along with Ser23 on helix α2 (**Fig. 2C and Suppl. Fig. S5**). Significant CSPs were also observed for Thr70 at the C-terminus of helix α4, and Leu41 located on the β2-strand (**Fig. 2D**). Both of these residues are positioned away from the main binding interface to DnaD DDBH2 and may indicate an allosteric effect induced in the presence of ssDNA as DnaD DDBH2 binds simultaneously ssDNA and the DnaA domain I.

Subsequently, a truncation of the DnaD DDBH2 domain (residues 129–196) was created to abolish DNA binding activity and investigate whether the C-terminally truncated DnaD maintained its DnaA binding activity without the region associated with binding nucleic acids. The truncated and unlabelled DnaD DDBH2 domain was titrated into ^15^N-labelled DnaA domain and subjected to the same CSP analysis. The results were fully consistent with those obtained for the full length domain with perturbations for the same nine key residues within DnaA domain I clearly evident. Significantly affected residues were located on helix α2 (Ser20, Ser23 and Thr26), helix α3 (Glu48, Phe49, Ala50 and Asp52) and helix α4 (His60) with an additional perturbation to Leu69 at the C-terminus of helix α4 also evident (**Fig. 2E, F and Suppl. Fig. S6**). These results confirm that the interaction between DnaA domain I and the DnaD DDBH2 domain is independent of the unstructured C-terminal region of DnaD (residues 206-215) essential to DNA binding, and largely unaffected by the binding of ssDNA.

### Identification of the DnaA interaction patch on ^15^N-labelled DnaD

We have previously described the assignment of the backbone of the DnaD DDBH2 domain using a similar methodology to that already described (**25**). The reverse titration of unlabelled DnaA domain I into a ^15^N-labelled DnaD DDBH2 domain (100 μM) up to an 8:1 excess now enabled us to map the complementary binding surface on DnaD DDBH2, using the same CSP methodology. A contiguous binding patch of nine residues on the DnaD DDBH2 domain was identified within the structured N-terminal region and involved two clusters of residues (**Fig. 3A, B and Suppl. Fig. S7**). The first of these, involved Leu129, Tyr130, Ile132, Phe133, Glu134, and Glu135 located on helix α1 and N-terminal region of the loop between helices α1 and α2. The second cluster involved Lys164, Glu169 and Val171 which were located throughout helix α3. Phe133 and Glu169 also showed CSP effects but are not surface exposed and hence perturbations may arise from small sidechain repacking associated with allosteric effects during the interaction. An averaged K_D_ = 665±251 μM calculated from the NMR titration data for a number of well-resolved residues (I132, V171, E135, Y130, E134) is fully consistent with the earlier estimate.

**Figure 3.**
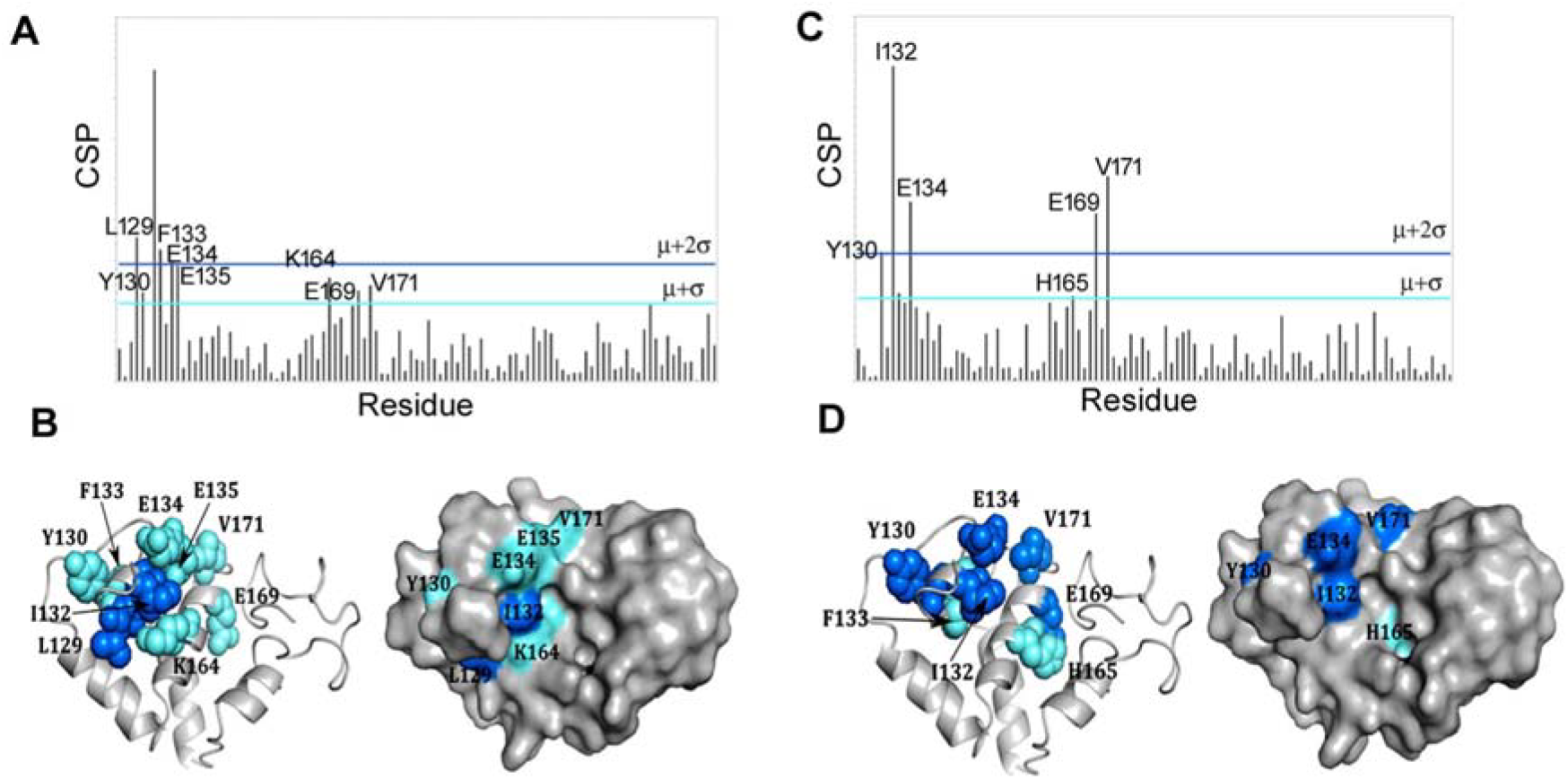
**A.** CSP analysis of the ^15^N-labelled DnaD DDBH2 domain titration with DnaA domain I. Threshold values of significance were calculated using μ = 0.00956 and σ = 0.00974. The μ + 2σ and μ + σ limits are highlighted in blue and cyan respectively. **B.** Residues exceeding the threshold limits are mapped onto the NMR structure of the DnaD DDBH2 domain (Marston et al., 2010). **C.** CSP analysis of the DnaD DDBH2 domain titration with DnaA domain I in the presence of ssDNA (800 μM). Threshold values of significance were calculated using μ = 0.00831 and σ = 0.00982. The μ + 2σ and μ + σ limits are highlighted in blue and cyan respectively. **D.** Residues exceeding the threshold limits are mapped onto the NMR structure of the DnaD DDBH2 domain.

The ternary complex of DnaA-DnaD interaction was further investigated using ssDNA (5’-GTTATTGCTC) in 8-fold excess. The addition of ssDNA to the experiment produced poorer quality HSQC spectra with weaker and broadened peaks displaying increased overlap between backbone amide residues (**Suppl. Fig. S8**). In particular, peaks corresponding to the unstructured C-terminal region of the domain were affected, and consequently certain amide resonances were excluded from the CSP analysis. Despite the poorer spectral resolution, two clusters of residues were observed; Tyr130, Ile132, Phe133 and Glu134 located on helix α1, and residues His165, Glu169 and Val171 in helix α3 (**Fig. 3C, D**). These are very similar to the residues identified in the equivalent CSP analysis in the absence of ssDNA (compare **Fig. 2C, D with 3C, D**).

### A model of the DnaA NTD-DnaD DDBH2 complex

Although the structure of *B. subtilis* DnaA domain I was solved by X-ray crystallography (Jameson *et al.*, 2014), the DnaD DDBH2 domain has a largely unstructured C-terminus rendering the DnaA domain I-DnaD DDBH2 complex unsuitable for structure determination using this technique. Instead, HADDOCK 2.2 was used to computationally model the interaction interface using the CSP effects as restraints (see Methods).

The DnaA domain I and DnaD DDBH2 structures and the two best-fit clusters to the NMR restraints are displayed in **Fig. 4**. Both clusters provide a DnaA binding interface distinct to the DNA binding patch within the DnaD DDBH2 domain. Of the 11 AIR restraints input for DnaA domain I, 8 and 9 were identified as interface residues within clusters 1 and 2 respectively, and of the 7 AIR restraints input for the DnaD DDBH2 domain, 5 were identified as interface residues for each cluster (**Suppl. Table S2**). The interaction surfaces of the individual clusters show overlap and the difference between the models can be accounted for by an approximated movement, for the DnaD DDBH2 domain, of 30 Å distance along an axis of rotation (**Fig. 4A, B**). The top clusters produced similar parameter scores which precluded the possibility of reliable discrimination between the two models. Moreover, we cannot eliminate the possibility that the two clusters represent distinct conformations of the complex in equilibrium in solution. Interestingly, neither cluster overlaps with the YxxxIxxxW motif and the F206-E215 region that have been shown previously to mediate the interaction of DnaD DDBH2 with ssDNA (**25**). This implies that DnaD is able to bind simultaneously the DnaA domain I and ssDNA.

**Figure 4.**
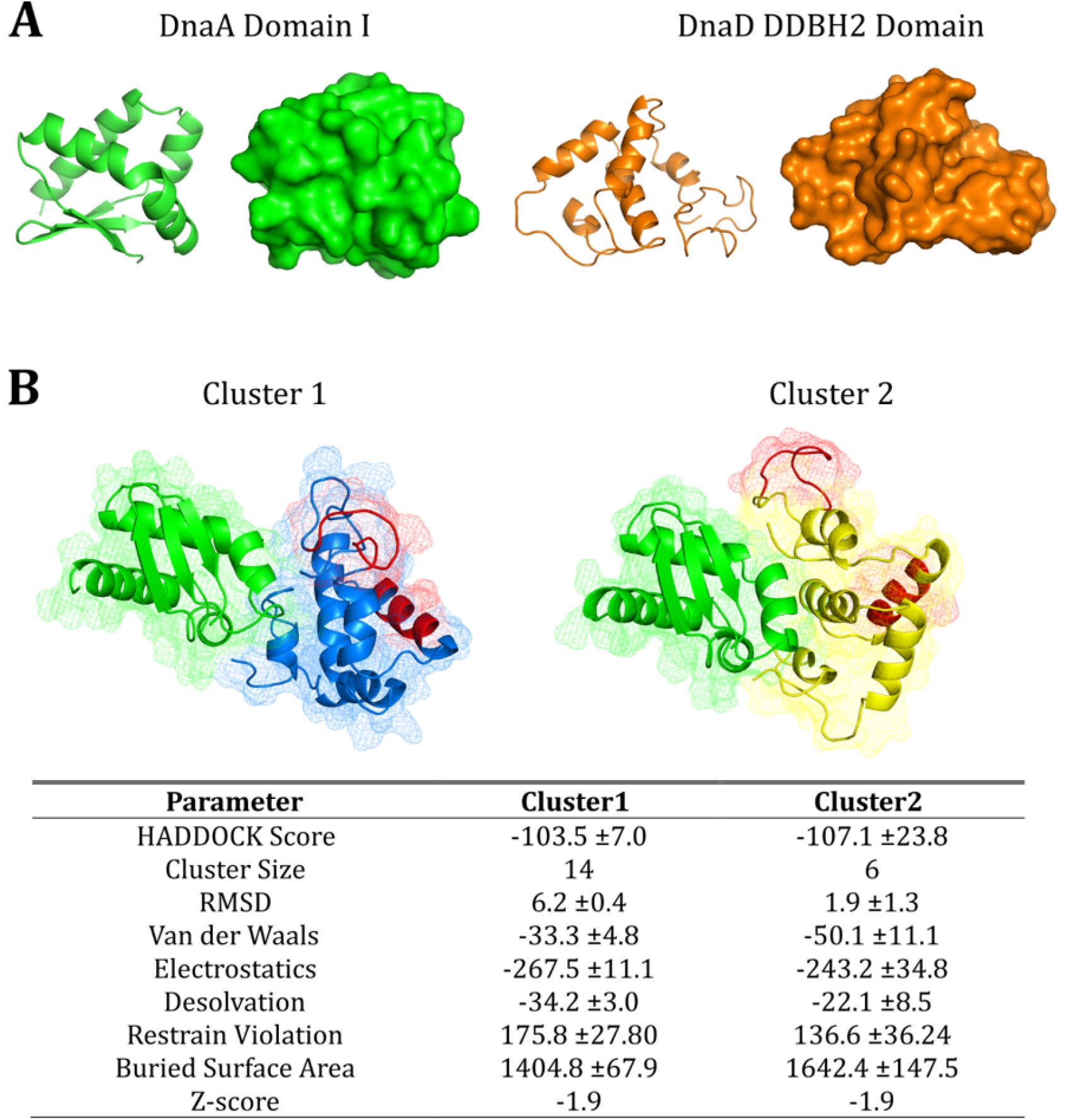
Predicted models of the DnaA domain I – DnaD DDBH2 domain interaction using HADDOCK restraint-driven docking. **A.** The structure of DnaA domain I (left, green) and the DnaD DDBH2 domain (right, orange) represented as cartoon and surface models. The structures were determined by X-ray crystallography (PDB – 4TPS) and NMR spectroscopy (Marston *et al.*, 2010) for DnaA domain I and the DnaD DDBH2 domain respectively. **B.** Top two clusters represented as mesh and cartoon, DnaA domain I is displayed in green and the DnaD DDBH2 domain is displayed in yellow (cluster 1) and blue (cluster 2), the DNA binding patch within the DDBH2 domain (YxxxIxxxW motif and F206-E215) is highlighted in red. The top clusters are displayed based on HADDOCK scoring and compliance with experimental restraints (from NMR chemical shift perturbation analysis).

### The interaction of DnaD with the DnaA-DNA filament induces filament untwisting

The abolition of BMOE-mediated cross-linking of higher order DnaA species within the DnaA-DNA filament in the presence of DnaD has been interpreted before as inhibition of filament formation by DnaD (**36**). However, an alternative interpretation of these data is that when DnaD interacts with the DnaA-DNA filament causing a conformational change that moves the N191C and A198C residues of adjacent DnaA molecules along the filament further away from each other preventing their physical crosslinking. The BMOE spacer is ~8 Å and any conformational change that moves N191C and A198C residues away from each other at a distance greater than 8 Å will prevent their BMOE-mediated cross-linking which could be mis-interpreted as inhibition of filament formation. In order to verify whether DnaD inhibits the formation of DnaA-DNA filaments we utilised single-molecule FRET experiments with N191C and A198C single mutant proteins fluorescently labelled with the FRET pair Cy3B and Atto647N. We hypothesized that if DnaD inhibits the formation of DnaA-DNA filaments there will be no detectable FRET in the presence of DnaD but if DnaD affects the conformation of the DnaA-DNA filaments then the FRET signal will be affected and FRET efficiency will be reduced but not abolished.

We used a model of a mini DnaA-DNA filament with four DnaA molecules reported before (**37**) to assess the interatomic distances of residues N191C and A198C within the filament and the feasibility of FRET experiments using these residues (**Suppl. Fig. S2**). Our modelling suggested that the interatomic distances are more appropriate for FRET with the N191C mutant protein and using a molar ratio of 1:1:2 DnaAN191CCy3B: DnaAN191CAtto647N:DnaAN191C should give us on average two labels per four DnaA molecules within the filament. In order to verify this, we constructed both DnaA N191C and A198C single mutant proteins and carried out FRET experiments to detect filament formation and compare FRET efficiencies between the two mutant proteins. FRET could be detected with both mutant proteins in the presence of ATP and DNA but it was somewhat better defined with DnaAN191C compared to DnaAA198C consistent with our modelling (**Fig. 5A and B)**. A small, high-FRET population could also be observed in the absence of ATP or DNA which is consistent with a tendency of DnaA molecules to weakly associate with each other (**Fig. 5C**).

**Figure 5.**
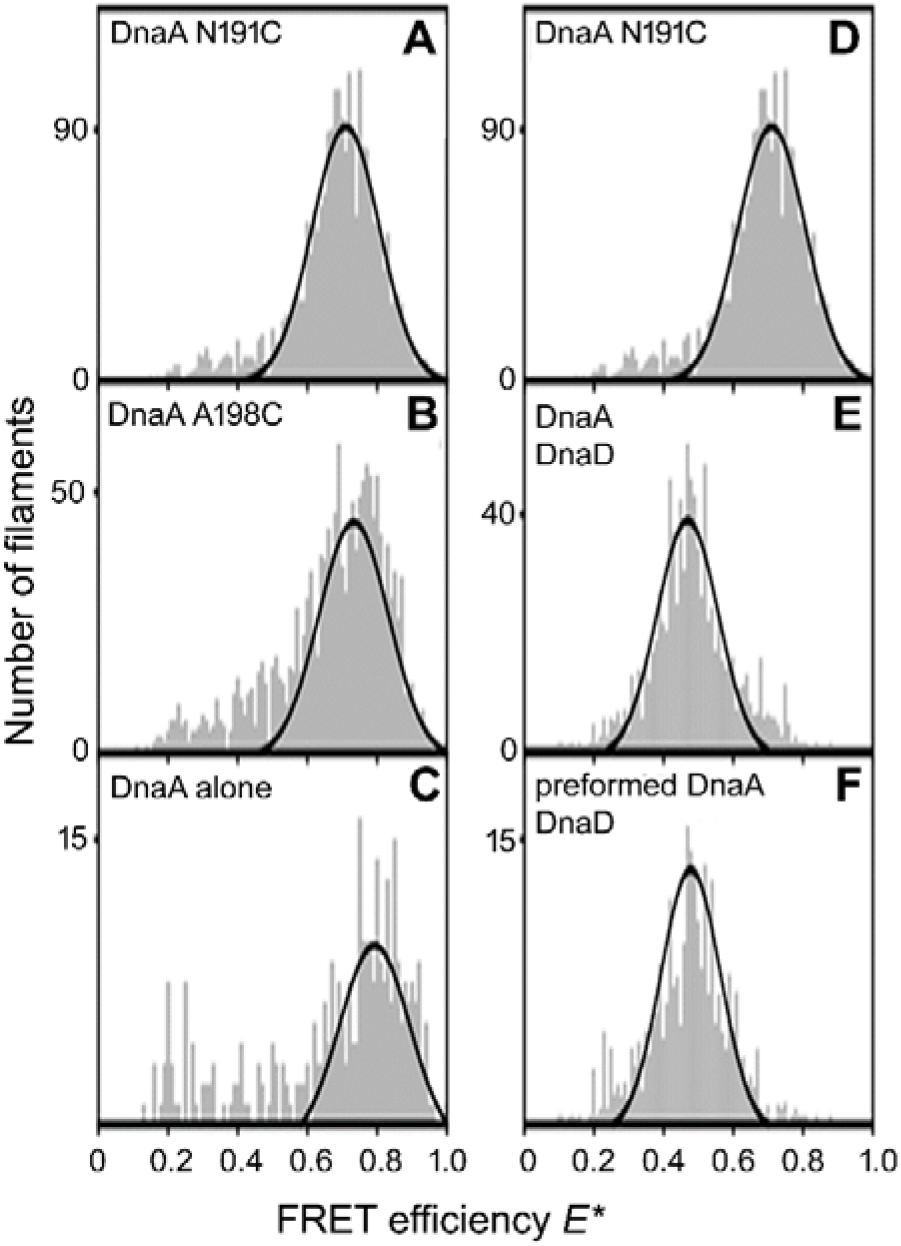
Histograms of the FRET efficiency *E\** determined from thousands of fluorescence bursts detected from labelled DnaA molecules diffusing through a confocal spot. (**A**) Filaments of DnaA labelled at residue N191C. (**B**) Filaments of DnaA labelled at positon A198C. (**C**) DnaA N191C in the absence of both DNA and ATP. (**D**) as in A. (**E**) DnaA filaments formed in the presence of DnaD. (**F**) DnaD added to preformed filaments of DnaA.

smFRET experiments were then carried out in the presence of DnaD (24 μM) added to the DNA either before or after the addition of DnaA in order to assess the effect of DnaD on the formation of DnaA-DNA filaments and also the effect of DnaD on pre-formed DnaA-DNA filaments (**Fig. 5D-F**). In the absence of DnaD high-FRET populations were detected with E*~0.7 (**Fig. 5D**) whereas in the presence of DnaD, either before or after the addition of DnaA, the FRET efficiency was shifted to E*~0.5 (**Fig. 5E, F**). These data are consistent with a DnaD-mediated conformational change (stretching and/or untwisting) of the DnaA-DNA filament that moves the fluorophores further apart from each other rather than complete inhibition of filament formation by DnaD. Furthermore, this DnaD-mediated untwisting can be induced on pre-formed DnaA-DNA filaments or during their formation in the presence of DnaD.

Further evidence that DnaD does not induce the disassembly of DnaA-DNA filaments was provided by electrophoretic mobility shift assays (EMSA) using a 120mer synthetic double stranded oligonucleotide containing the four DnaA boxes within the half origin between *dnaA* and *dnaN* (see **Suppl. Fig. S10**).

### DnaD-mediated stretching/untwisting of DnaA-DNA filaments does not require DnaD binding to DNA

The structure of the complex between DnaA domain I and the DnaD DDBH2 revealed that the DNA-interacting region of DDBH2 is distinct and does not overlap with the DnaA-interacting patch of DDBH2. Therefore, DnaD can potentially interact simultaneously with the domain I of DnaA and with the DNA that is wrapped around the outside of the DnaA-DNA filament. It is not clear whether both interactions of the DnaD DDBH2 with the domain I of DnaA and DNA are required to untwist the filament. In order to investigate this we used DnaD196 which is a truncated version of DnaD lacking the C-terminal region residues 197-232 abolishing its ability to bind to DNA (**25**). We carried out comparative experiments with full length DnaD and DnaD196 to compare their effects on DnaA-DNA filaments. Both full length DnaD and DnaD196 appeared to have similar effects on the filaments shifting the FRET efficiency from *E\**~0.7 in their absence to *E\**~0.5 in their presence. This suggests that both DnaD and DnaD196 untwist the filament to a similar extend (**Fig. 6A-C**) and therefore DnaD binding to DNA around the outside of the filament is not required for this DnaD-mediated conformational change to occur.

**Figure 6.**
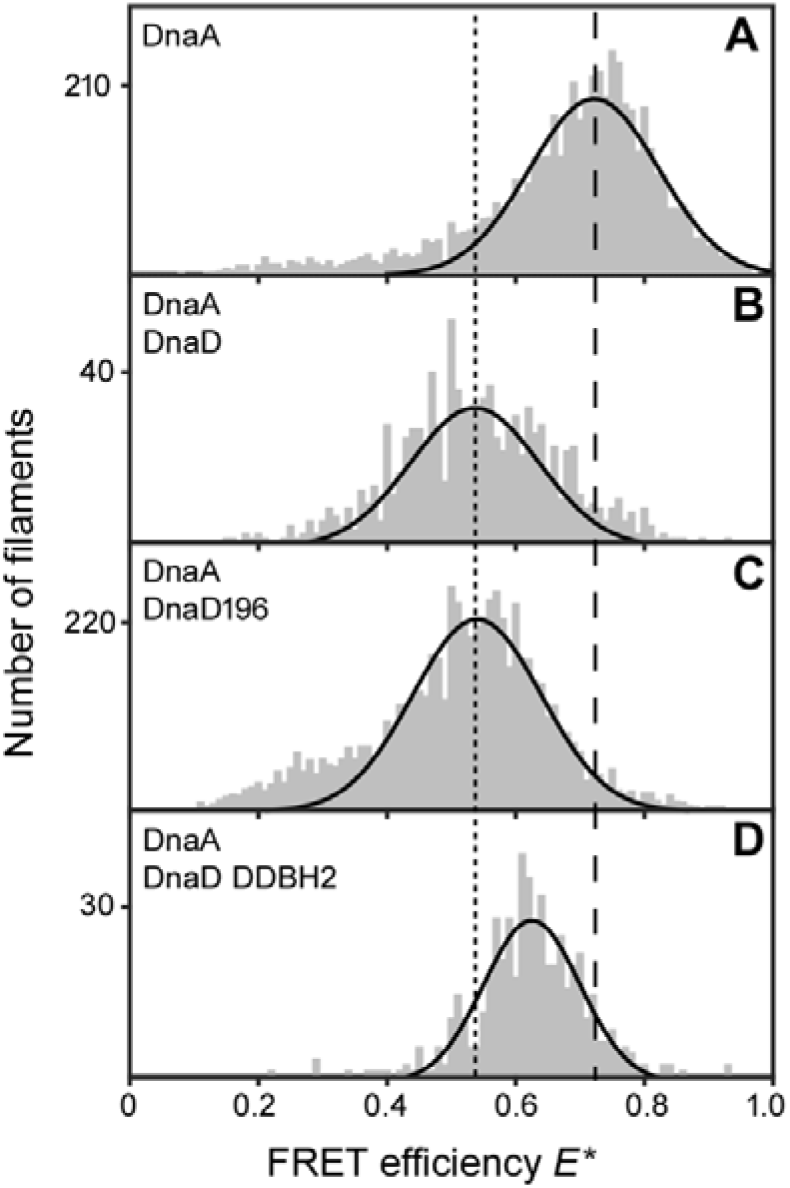
Histograms showing the effect of DnaD on DnaA inter-subunit distances as determined by smFRET. (**A**) DnaA filaments alone. (**B**) DnaA filaments incubated with full-length DnaD (24 uM). (**C)** DnaA filaments incubated with DnaD196 (24 uM). (**D**) DnaA filaments incubated with DnaD DDBH2 (24 uM). Data were fit with single-Gaussian distributions (solid black lines). The dashed line marks the mean E* value for the DnaA filament alone. The dotted line marks the mean E* value for the DnaA filament incubated with full length DnaD.

Interestingly, the DnaD DDBH2 on its own, which retains its ability to interact with the domain I of DnaA and DNA but lacks the scaffold-forming N-terminal domain (DDBH1) of DnaD (**28, 29**), also affected the conformation of the DnaA-DNA filament since the FRET efficiency shifted from *E\**~0.7 in its absence to *E\**~0.6 in its presence (compare **Fig. 6A and D**). This is somewhat different than the *E\**~0.5-value apparent in the presence of full length DnaD or DnaD196 (compare **Fig. 6B, C with D**), suggesting that DDBH2 can also induce a conformational change of the DnaA-DNA filament but smaller than that observed in the presence of full-length DnaD or DnaD196.

## Discussion

### Both the DDBH1 and DDBH2 domains of DnaD interact with DnaA domain I and DnaD can interact simultaneously with DnaA and DNA

A recent bacterial two hybrid study suggested that the DDBH1 domain of DnaD interacts with the DnaA domain I while the DDBH2 does not (**34**) contradicting an earlier yeast two hybrid study which showed that residues 133-140, at the N-terminal end of the DDBH2 domain, are crucial for the DnaD-DnaA interaction (**33**). Our data reveal that both DDBH1 and DDBH2 interact with the DnaA domain I. The interaction of the DDBH1 with the DnaA domain I appears to be stronger than that of DDBH2 and was not amenable to straight forward NMR investigations, as the presence of intermediate exchange resulted in loss of signal.

However, the interaction of DDBH2 with the DnaA domain I was studied in detail by NMR and HADDOCK modelling using the experimental restraints from the CSP analysis of the NMR data and revealed two possible overlapping, but distinct, families of conformations (**Fig. 4**). In addition, the importance of the DDBH1 WH (Wing Helix) motif for the DnaD-DnaA interaction suggest that this interaction has an extensive interface encompassing both the DDBH1 and DDBH2 domains with two slightly different conformations. In the context of the native DnaD protein which forms a core tetramer mediated via DDBH1 interactions (**28**), the interaction with DnaA will involve both the core DDBH1 tetramer and the individual DDBH2 domains likely projecting out of the core DDBH1 tetramer. Interestingly, the interactions of DnaD with DnaA and ssDNA appear to involve separate patches that do not overlap with each other, suggesting that DnaD can potentially bind simultaneously to both ssDNA and DnaA.

### The DDBH1 and DDBH2 interaction patches on DnaA domain I overlap

The DnaA-interaction patch of DDBH2 contained residues (F133, E134 and E135) within the amino acid region 133-140 that was previously identified as essential by yeast two hybrid analysis (**33**). This region appears to be part of a wider interaction patch extending to residues L129, Y130, K164, E169, V171 and H165 (**Fig. 3**). In the recent bacterial two hybrid analysis residues F49, W27 and T26 of DnaA domain I were identified as important for the interaction with the DDBH1 (**34**). Our data revealed a more extended network of residues in the DnaA domain I that are involved in the interaction with the DDBH2 encompassing residues K17, S20, S23, F49, A50, D52, E55, W53 and H60 (**Fig. 2**), with F49 identified by both studies. Furthermore, residue T26 was also identified by our study in the interaction of the truncated DDBH2 (residues 129-196) with the DnaA domain I (**Fig. 2E**). Therefore, it appears that the interactions of the DDBH1 and DDBH2 domains with the DnaA domain I involve overlapping patches.

### DnaD-mediated stretching/untwisting of the DnaA-DNA filament

Critically, the interaction of full-length DnaD with DnaA appears to be cryptic and could not be detected by bacterial two hybrid (**34**) suggesting that DnaD conformational changes are required to render it competent to bind DnaA. However, we and others were able to detect an interaction of full-length DnaD with DnaA-DNA filaments using single molecule FRET and/or a BMOE-mediated cross-linking technique (this study and **36**). One possibility could be that, in the context of DnaA-DNA filaments, DnaD conformational changes may result from its initial interaction with the DNA which then render it competent to simultaneously bind to DnaA thus inducing filament untwisting. However, our single molecule FRET studies revealed that DnaD196, which still forms tetramers but lacks the C-terminal residues 197-232 and is incompetent in binding DNA (**25**), was able to untwist the DnaA-DNA helix equally well as the full-length DnaD (**Fig. 6A-C**), suggesting that DnaD binding to DNA is not required to untwist the filament. Therefore, the filament untwisting is likely the result of a DnaD-induced conformational change of the DnaA filament which stretches and untwists the DNA wrapped around the outside of the DnaA filament.

Interestingly, the DDBH2 domain was also able to untwist the filament but to a lesser extent than full-length DnaD (**Fig. 6A-D**), suggesting that either the DDBH1 domain also contributes to the DnaA filament untwisting or a DnaD tetramer is required to fully untwist the filament. Full-length tetrameric DnaD by itself has DNA untwisting activity (**30, 31**), extending B-form duplex DNA from 10.5 bp per helical turn to 16 bp per turn (**32**). Single-molecule atomic force spectroscopy studies revealed that the DDBH2 domain by itself can also stretch/untwist DNA (**31**) which is also consistent with our single molecule FRET studies. Collectively our data suggest that one of the functions of DnaD during replication initiation may be to work cooperatively with DnaA in order to stretch/untwist the filament and help to locally melt the *oriC*.

### DnaD and SirA are competitive regulators of DnaA function

The DnaA domain I interacts with the replicative helicase and a number of client proteins involved in the regulation of replication initiation (**16-21**) and as such it appears to be a regulatory interaction hub. DnaD can now be added to the list of client proteins that interact with this regulatory hub.

Our data show the DnaA-DnaD binding interface of the DnaA domain I overlaps with the DnaA-SirA interface determined by X-ray crystallography (**15**). This directly confirms a recent bacterial two hybrid study which has also revealed this overlap (**34**). SirA is a negative regulator of DNA replication, it inhibits initiation of replication in diploid cells committed to sporulation. The DnaA-SirA interaction surface is α-helical with both proteins using polar side chains packing against each other within a predominantly hydrophobic interface. Both DnaD and SirA interact with domain I of DnaA and their interaction patches highly overlap with each other (**Fig. 7**). SirA binding could sterically hinder the DnaA-DnaD interaction to prevent re-initiation of DNA replication in *B. subtilis* cells committed to sporulation. It has been suggested before that SirA may inhibit the DnaA-DnaD interaction arresting assembly of the initiation complex (**15, 38**). This is consistent with our data and suggests that DnaD has a positive role during replication initiation as opposed to the negative regulatory role suggest by others (**39**). Therefore, SirA and DnaD achieve opposing regulatory functions via interactions with the same structural site on DnaA.

**Figure 7.**
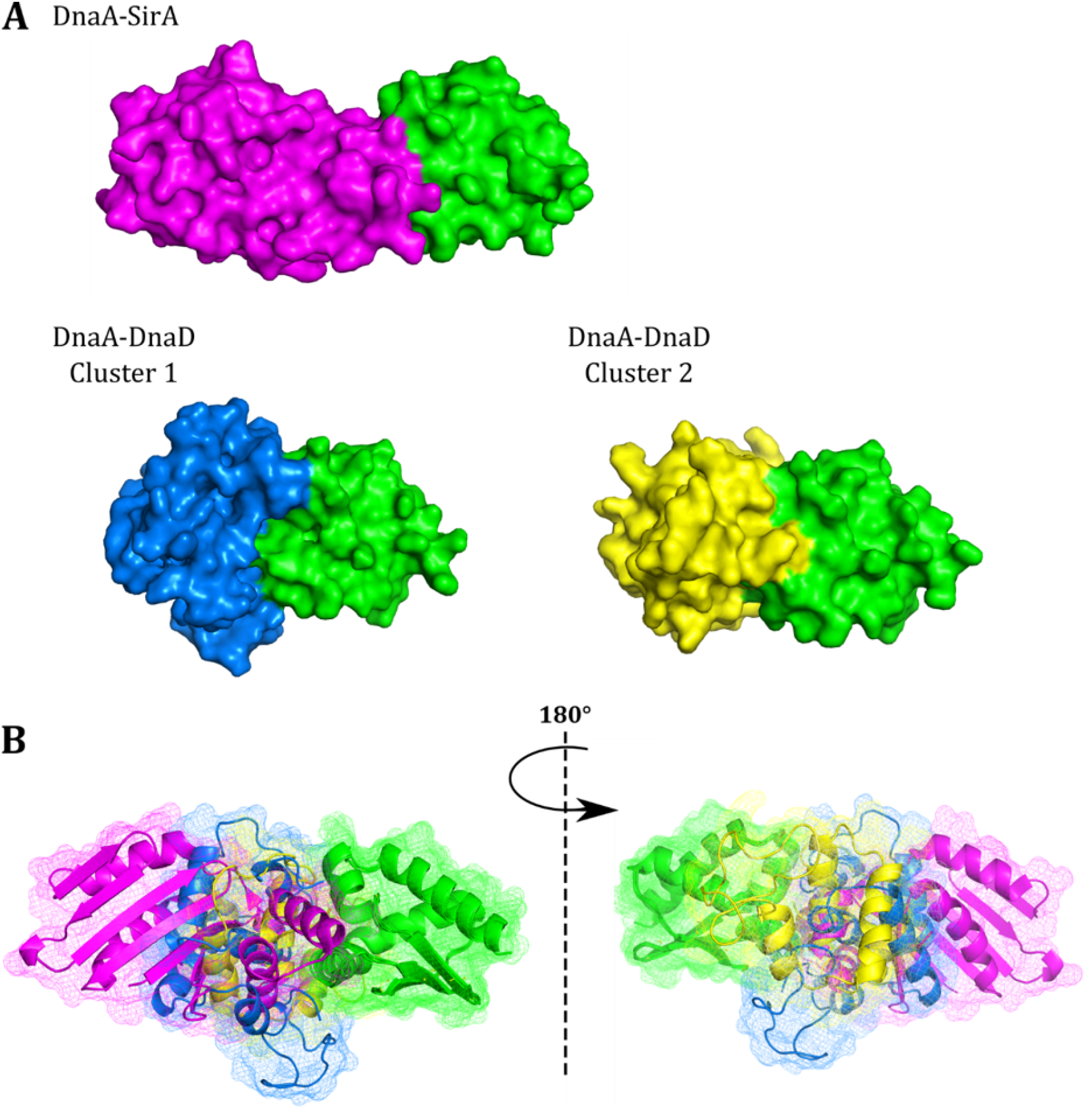
The DnaA-DnaD interface overlaps with the DnaA-SirA binding interface. **A.** The structure of DnaA-SirA (left), and predicted models of the DnaA-DnaD interaction Cluster 1 (centre) and Cluster 2 (right) represented as surface models. DnaA domain I is displayed in green, SirA displayed in purple and the DnaD DDBH2 domain is displayed in blue and yellow for cluster 1 and cluster 2 respectively. The structure of the DnaA-SirA binding interface was determined by X-ray crystallography (Jameson *et al.*, 2014), (PDB – 4TPS) **B.** Overlay of the DnaA-SirA complex with the two predicted DnaA-DnaD complex models displayed as cartoon and mesh representation. Individual domains are coloured as described for A. There is considerable overlap between the SirA binding interface and the DnaD binding interface suggesting that SirA binding could sterically hinder the DnaA-DnaD interaction to prevent re-initiation of DNA replication in B. subtilis cells committed to sporulation.

## Methods

### Cloning, Expression and Purification

Domain I (residues 1-81) of the *B. subtilis* DnaA protein was cloned into the pET28b+ vector for expression as an N-terminal histidine-tagged protein. The pET28b+ construct to express and purify the DDBH2 domain (residues 129-232) of the *B. subtilis* DnaD protein was supplied by Dr Jeremy Craven (**25**) and DnaD DDBH1 was produced as described before (**28**). A truncated version of the DnaD DDBH2 domain (residues 129-196) was produced using Q5 site directed mutagenesis. This method was also used to produce all of the DnaA domain I mutants. XL1-Blue electrocompetent cells were used for plasmid storage and CaCl_2_ competent BL21(DE3) cells were used for expression of these constructs. BL21(DE3) cells were grown in M9 minimal media supplemented with ^15^N ammonium chloride and/or ^13^C glucose, for use in NMR spectroscopy, where isotopic labelling was required. Soluble protein was obtained by induction with IPTG (1 mM) at mid-log growth phase (OD_595_ 0.6-0.8) followed by expression for 16 hr at 30°C. Soluble protein was obtained by induction with IPTG (1 mM) at mid-log growth phase (OD_595_ 0.6-0.8) followed by expression for 4 hr at 30°C or 16 hr at 20°C for DnaA and DnaD constructs, respectively. Sonication (10 microns, 4 min) followed by centrifugation (35,000 *g*, 30 min) was used to obtain the soluble fraction of the cell lysate. Ni-affinity chromatography (50 mM potassium phosphate, 0.5 M NaCl, pH 8 with a gradient to 0.25 M imidazole) was used as a first stage of purification followed by thrombin cleavage of the His-tag (16 hours at room temperature in 20 mM Tris-HCl pH 8.4, 150 mM NaCl, 2.5 mM CaCl_2_) leaving behind a residual N-terminal Gly-Ser-His-tag. Ni-affinity chromatography was repeated for further separation of the His-tagged from the un-tagged protein. This was followed by size exclusion chromatography using a Superdex 75 26/60 column (50 mM potassium phosphate, 100 mM NaCl, pH 7.4) as the final purification stage. DnaA constructs were desalted into pure MilliQ-water and lyophilised for storage whereas DnaD constructs were frozen in solution.

### NMR Spectroscopy

NMR experiments were recorded at 25°C on a Bruker 800 MHz Avance III spectrometer with a QCI cryoprobe. Data acquisition and processing were carried out using Topspin 3.1.b.53 software and were further compared within the CCPNMR software.. 3D NMR experiments used ^13^C^15^N labelled DnaA domain I at 600 μM (50 mM potassium phosphate pH 7.4, 50 mM NaCl, 5% v/v D_2_O, 0.02% w/v NaN_3_). Backbone resonances were assigned using HNCO, (HCα)CONH, CβCαNH and CβCα(CO)NH experiments with watergate suppression.

Backbone assignment was facilitated through selective unlabelling experiments to resolve some spectral ambiguities, signal overlap and exclude particular reside types from the ^15^N HSQC spectra. This approach was successful for unlabelling of lysine, arginine, asparagine, and histidine residues. Similarly, to aid assignment, single point mutations of a number of surface exposed residues (S75A, T26A, G38A, S56A, E68N, S23A) were used to locally perturb signals in the ^15^N HSQC experiment. The combination of sequentially linking amino acids from the 3D spectra and these additional approaches allowed 100% assignment of non-propyl residues 1-81 of DnaA domain I ^15^N HSQC spectrum. Three residues remained unassigned corresponding to a short, flexible leader sequence Gly-Ser-His resulting from the thrombin cleavage of the N-terminal His-tag.

Chemical shift perturbations (CSPs) where calculated based upon a weighting of ^1^H and ^15^N shifts according to the equation CSP = √ ½ [δ^2^H + (0.14 x δ^2^_N_)]. Significant CSP effects were determined using the standard deviation (σ) from the mean (μ) and residues with CSP values above ‘μ + σ’ and ‘μ + 2σ’ (**40**) were used for interaction surface mapping and structural modelling. NMR titration experiments used ^15^N isotopically labelled DnaA domain I or ^15^N DnaD DDBH2 domain at 100 μM concentration (50 mM potassium phosphate pH 7.4, 100 mM NaCl, 5% v/v D_2_O, 0.02% w/v NaN_3_). ^15^N HSQC spectra were collected at 0.5 molar ratio intervals from 0:1 to 8:1 excess of DDBH2 or domain I.

For the HSQC titration of DnaD DDBH1 into DnaA domain I, a stock solution of ^15^N isotopically labelled DnaA domain I (100 μM) in 50 mM potassium phosphate pH 7.4, 100 mM NaCl, 10% v/v D_2_O, 0.02% w/v NaN_3_. A DnaD DDBH1 stock solution (400 μM) was also prepared in the same buffer. Aliquots of the DnaD DDBH1 solution were then titrated into the DnaA domain I solution to yield solutions at various stochiometric ratios.

### Structural model

HADDOCK 2.2 (High Ambiguity Driven protein-protein Docking) accessible via the WeNMR (**41**) server was used to computationally model the DnaA domain I/DnaD DDBH2 interaction interface using the experimental restraints from the CSP analysis of the NMR data. The HADDOCK docking protocol consists of rigid-body docking, semi-flexible refinement, and final refinement in explicit solvent. Ambiguous interaction restraints (AIRs) were generated prior to running HADDOCK, these used experimental data, such as NMR CSP data, to define ‘active’ residues (experimentally derived and solvent accessible), and ‘passive’ residues (solvent accessible residues neighbouring active residues). To account for errors in the definition of active and passive residues, HADDOCK allows the random deletion of a fraction of the restraints for each docking run. Subsequently, multiple runs using the same AIR input were used to prevent bias. The HADDOCK score given to output models was a weighted sum of intermolecular electrostatics, van der Waals, desolvation, and AIR restraints. A z-score was also given which represents how many standard deviations the HADDOCK score of a given cluster is away from the mean of all clusters. Multiple HADDOCK runs were undertaken using the AIR restraints shown in the **Suppl. Table S1**. Initially, restraint inputs that were varied to confirm output models were not purely driven by energy forces (electrostatics, hydrophobics, Van der Waals). Using identical AIR inputs for 10 HADDOCK runs, 162 structures in 13 clusters were generated. The clusters were then analysed to select the best models to fit with experimental restraints provided by the CSP data.

### Formation of DnaA-DNA filaments

DnaA-DNA filaments were formed using the DnaA(N191C/A198C) mutant protein and 819 bp DNA fragment containing the half origin with the DUE and four DnaA-boxes between the *dnaA* and *dnaN* genes (**Suppl. Fig. S2 and S3**) as described before (**36**). Briefly, 1.5 μM DnaA(N191C/A198C) was incubated with 3 nM DNA, representing 500:1 molar ratio of DnaA:DNA at 37°C for 15 mins, in 25 mM HEPES pH 7.6, 200 mM NaCl, 100 mM potassium glutamate, 10 mM MgCl_2_ and 2 mM ATP. BMOE (2 mM) was then added and the mixture was incubated at 37°C for an additional 5 mins before the addition of cysteine (50 mM) to quench the cross-linking reaction for 5 mins. Proteins were then resolved through SDS-PAGE and visualised via Coomassie staining. Experiments were carried out in the presence and absence of DnaA domain I, as indicated.

### Fluorescent Labelling of DnaA

Single cysteine DnaA mutants N191C and A198C were labelled with each of the Atto647N-maleimide and Cy3B-maleimide dyes. Atto647N-maleimide or Cy3B-maleimide (10 mM in DMSO) was added dropwise at 10 x molar excess to proteins DnaA^N191C^ and DnaA^N198C^ in 50 mM Tris pH 7.5, 100 mM NaCl. The reaction mixture was flushed with nitrogen and the reaction was left to proceed overnight at 4**°**C with rotation mixing, prior to being quenched with 1 mM DTT. The reactive dyes were shielded from direct light throughout the labelling procedure. Atto647N-labelled and Cy3B labelled DnaA^N191C^ and DnaA^A19C^ were seperated from excess fluorescent dyes by extensive dialysis overnight at 4**°**C (into an appropriate buffer) was repeated until no further excess dye molecules were present in the exchanged buffer.

### Single-molecule FRET experiments

Single-molecule FRET measurements were performed at room temperature using a home-built confocal microscope (as previously described **42**). Briefly, the microscope operated with 20 kHz alternating-laser excitation between a 532-nm (Samba, Cobolt, operated at 240 μW) and a 638-nm laser (Cube, Coherent, operated at 60 μW), coupled to a 60x, 1.35 numerical aperture (NA), UPLSAPO 60XO objective (Olympus). Emitted photons were spectrally filtered and detected by two avalanche photo diodes (SPCM-AQRH-14, Perkin Elmer). The alternating laser excitation allows filtering for correctly labelled species bearing an active donor and an active acceptor (**43**). After filtering each fluorescent burst for the correct labelling stoichiometry, we calculated the apparent FRET efficiency E* for each burst as E*=DA/(DD+DA), where DA is the number of photons in the red detection channel after green excitation and DD the number of photons in green detection channel after green excitation.

Filaments were formed as described above, with a total of 1.5 μM DnaA (ration of 1:1:2 Cy3B-labelled: Atto647N labelled: un-labelled) incubated with 3 nM DNA ± DnaD (24 μM), at 37°C for 15 mins, in 25 mM HEPES pH 7.6, 200 mM NaCl, 100 mM potassium glutamate, 10 mM MgCl_2_ and 2 mM ATP. Measurements were taken in ‘Imaging buffer’, consisting of 40□mM 4-(2-hydroxyethyl)-1-piperazineethanesulfonic acid (HEPES)-NaOH, pH 7.3, 10□mM MgCl_2_, 1□mM DTT, 100□μg□ml−1 bovine serum albumin, 5% (vol/vol) glycerol, 1□mM mercaptoethylamine, containing ~100 pM DnaA.

### Oligonucleotides

Oligonucleotides used for protein cloning are shown in **Suppl. Fig. S11.** Oligonucleotides used for mutagenesis of DnaA domain I during NMR structure determination are shown in **Suppl. Fig. S12**. Protein mutagenesis was carried out with the New England Biolabs Q5 site directed mutagenesis kit, as described by the manufacturer.

## Acknowledgments

We thank Dr. Jeremy Craven (University of Sheffield) for provision of materials, assignment of NMR data and useful discussions. This work was supported by a BBSRC grant (BB/R013357/1) to P.S.. M.P. is a PhD student partially funded by a Vice Chancellor’s Excellence Award at the University of Nottingham. E.M. was supported by a PhD studentship funded by the School of Chemistry, University of Nottingham.

